# Aperiodic component of EEG power spectrum and cognitive performance in aging: the role of education

**DOI:** 10.1101/2023.10.05.560988

**Authors:** Sonia Montemurro, Daniel Borek, Daniele Marinazzo, Sara Zago, Fabio Masina, Ettore Napoli, Nicola Filippini, Giorgio Arcara

## Abstract

Aging is associated with changes in the oscillatory -periodic-brain activity in the alpha band (8-12 Hz), as measured with resting-state EEG (rsEEG); it is characterized by a significantly lower alpha frequency and power. Aging influences the aperiodic component of the power spectrum: at a higher age the slope flattens, which is related with lower cognitive efficiency. It is not known whether education, a cognitive reserve proxy recognized for its modulatory role on cognition, influences such relationship.

N=179 healthy participants of the LEMON dataset (Babayan et al., 2019) were grouped based on age and education: young adults with high education and older adults with high and low education. Eyes-closed rsEEG power spectrum was parametrized at the occipital level.

Lower IAPF, exponent, and offset in older adults were shown, compared to younger adults. Visual attention and working memory were differently predicted by the aperiodic component across education: in older adults with high education, higher exponent predicted slower processing speed and less working memory capacity, with an opposite trend in those with lower education.

Further investigation is needed; the study shows the potential modulatory role of education in the relationship between the aperiodic component of the EEG power spectrum and aging cognition.

## 1. INTRODUCTION

As we age, many changes occur in individuals’ behavior and cognition, worsening in memory ^1^, attention span ^2^, executive functions ^3^, and processing speed ^4^, reflecting just a few of the consequences of the natural aging process. These behavioral changes are accompanied by (and associated with) changes in the brain’s structural anatomy ^5,6^, metabolism ^7^ and functionality ^8^ which produce a significant effect on its neurophysiological activity ^9^. Electroencephalogram (EEG) studies on aging have shown changes in neural oscillatory activity, especially in the alpha band (8-12 Hz) ^10–12^. Researchers have reported that older adults display slower alpha oscillatory activity and lower alpha power than their younger counterparts ^13–15^. Moreover, individual alpha peak frequency (IAPF), i.e., the frequency where EEG activity exhibits the maximum power in the alpha range, tends to decrease from adulthood to midlife ^11,12^. In most studies, EEG activity in specific frequency bands has been traditionally measured as the average of the power in the frequency bands of interest as calculated from the power spectrum ^16^. However, this approach (and subsequent interpretation of the results) has been challenged by a renewed interest in the non-oscillatory, aperiodic component of the EEG signal.

The aperiodic component does not necessarily entail specific frequency bands; it exhibits a 1/f-like distribution in the log-log space of a Power Spectrum Density (PSD), meaning its power exponentially decreases as frequency increases.

Aperiodic activity can be parametrized by values of the exponent, which describe the negativity of the power spectrum slope and the offset, the broadband shift of power across frequencies ^16^. Importantly, change in the spectrum’s aperiodic component may occur without changes in the oscillatory components and may affect the power values calculated for each frequency. For example, a change in aperiodic slope may influence the power values in the alpha band as calculated from the power spectrum, when no change in alpha’s oscillatory component occurred. This may lead to spurious results and wrong interpretations and highlight the importance of taking into account the aperiodic component when interpreting power spectrum data ^16,17^.

For a long time, researchers ignored the aperiodic component of brain activity, considering it to be merely background noise; however, recently, this component has gained attention not only for its methodological impact but also because of its potential role as a marker of neurophysiological mechanisms underlying cognition. For instance, the aperiodic offset has been associated with neuronal population spiking ^18,19^ and blood-oxygen-level-dependent (BOLD) signal of functional magnetic resonance (fMRI) ^20^. Meanwhile, the exponent has been linked to changes in cortical excitation/inhibition balance ^21^, which means that increased power in higher frequency (associated with increased excitation at the cerebral level) produced a flattening of the PSD; thus a diminished exponent; the exponent has also been related to local neural tissue properties ^22,23^. In light of the investigations mentioned above, narrowly associating 1/f characteristics with specific cerebral function, alternative explanations should be acknowledged. A possible interpretation of the functional meaning of the aperiodic component posits that this variable indicates an underlying self-organized critical system ^24^. Within this theoretical framework, the critical system is considered as the optimal state of brain’s functionality ^25^ where large-scale reorganization occurs quickly in response to stimuli, reflecting the brain’s adaptation to changes in the endogenous and exogenous environment ^26^. Aperiodic activity may arise from spatial integration of asynchronous spiking of neural populations ^27^. Therefore, a reduction of the spectral exponent suggests functional neural decoupling.

In the context of aging, a first study from ^28^ showed that the aperiodic slope of EEG and electrocorticography (ECoG) spectra flattened in a group of older people compared with a younger one, with decreased power between 8 to 14 Hz and increased power between 14 and 25 Hz. The changes in EEG spectral slope were also associated with age differences in working memory performance. Interestingly, the aperiodic exponent appeared to mediate this relationship, suggesting that the slope effect alone could account for behavioral differences between older and young adults. Voytek and colleagues explained these results with the “Neural Noise Hypothesis”. Initially proposed by ^28^, this theory suggests that as people age, there is an increase in spontaneous desynchronized neural activity, resulting in a decreased fidelity of neural communication and a flatter power spectrum. Recent studies have replicated Voytek findings and added new insights to the relationship between aging and aperiodic spectral components, highlighting their impact on cognitive performance. Flatter slopes in older adults were linked to poorer performance during spatial attention tasks ^29,30^ and short-term memory tasks ^31^. Recent work from ^32^ has associated the flattening of the aperiodic slope in frontal regions with worse performance on tasks involving processing speed and executive functions. Moreover, ^29^ measured changes of the aperiodic spectra at the baseline period in younger and older adults. They examined to what extent 1/f like exponent was related to alpha trial-by-trial consistency, during a spatial discrimination task. The authors considered these signal properties as measures of “neural noise”. Interestingly, they found that older adults with the highest baseline noise levels also had the worse alpha trial-by-trial consistency, suggesting that age-related increases in baseline noise might diminish sensory processing and cognitive performance.

Understanding the impact of neural noise could unveil new perspectives in capturing the effects of aging on neurophysiological functioning. Critically, studies on the aperiodic component of the spectrum and its functional role have neglected the possible modulatory role of “cognitive reserve” in modulating the effects on cognition in aging. Cognitive reserve is a concept proposed by Stern ^33,34^, and it describes the capacity of the brain to actively cope with age-related or pathological changes by employing pre-existing resources. Such reserve accumulates since early life through exposure to cognitively stimulating life ^35–37^. One of the most widely recognized variables associated with the level of cognitive reserve is education, typically operationalized as a numerical variable, indicating years of successful education or as an educational level operationalized as an ordered factor (e.g., high school, University etc.). Education has been largely used to examine the aging population’s potential compensatory processes ^38,39^. While education modulates cognitive performance (i.e., the higher the education, the better cognitive performance), its impact on the intricate relationship between neural activity in the aging brain and cognitive capacity remains uncertain. Aperiodic activity, conceived as a possible pattern of neural communication impairment, is associated with a decline in cognitive performance in older adults. Critically, although education can be expected to favor the preservation/maintenance of neurobehavioral functionality in aging ^40^, how this is reflected at the level of aperiodic brain activity is not clear yet.

With reference to the most recently updated framework targeting aging compensatory mechanisms, a specific set of variables are combined in this work, to investigate: i) the possible age-related neural patterns (i.e., aperiodic values), ii) age-related cognitive differences (test scores), iii) and education as a variable expected to moderate the association between i) and ii) ^34^. Older adults with higher education were expected to preserve a more youth-like profile as compared to older adults with lower education.

## 3. RESULTS

### 3.1. Descriptive Analyses

Young adults performed significantly better on all cognitive tasks than older adults (Table S1 and Figure S1). IAPF, exponent, and offset values were shown in young adults compared to older adults (Figure 3 and S1). Older adults with different education did not differ on most cognitive tasks except for the working memory one, where older adults with high education performed better than older adults with lower education and more similarly to the younger adults (Table 1).

**Figure 1.**
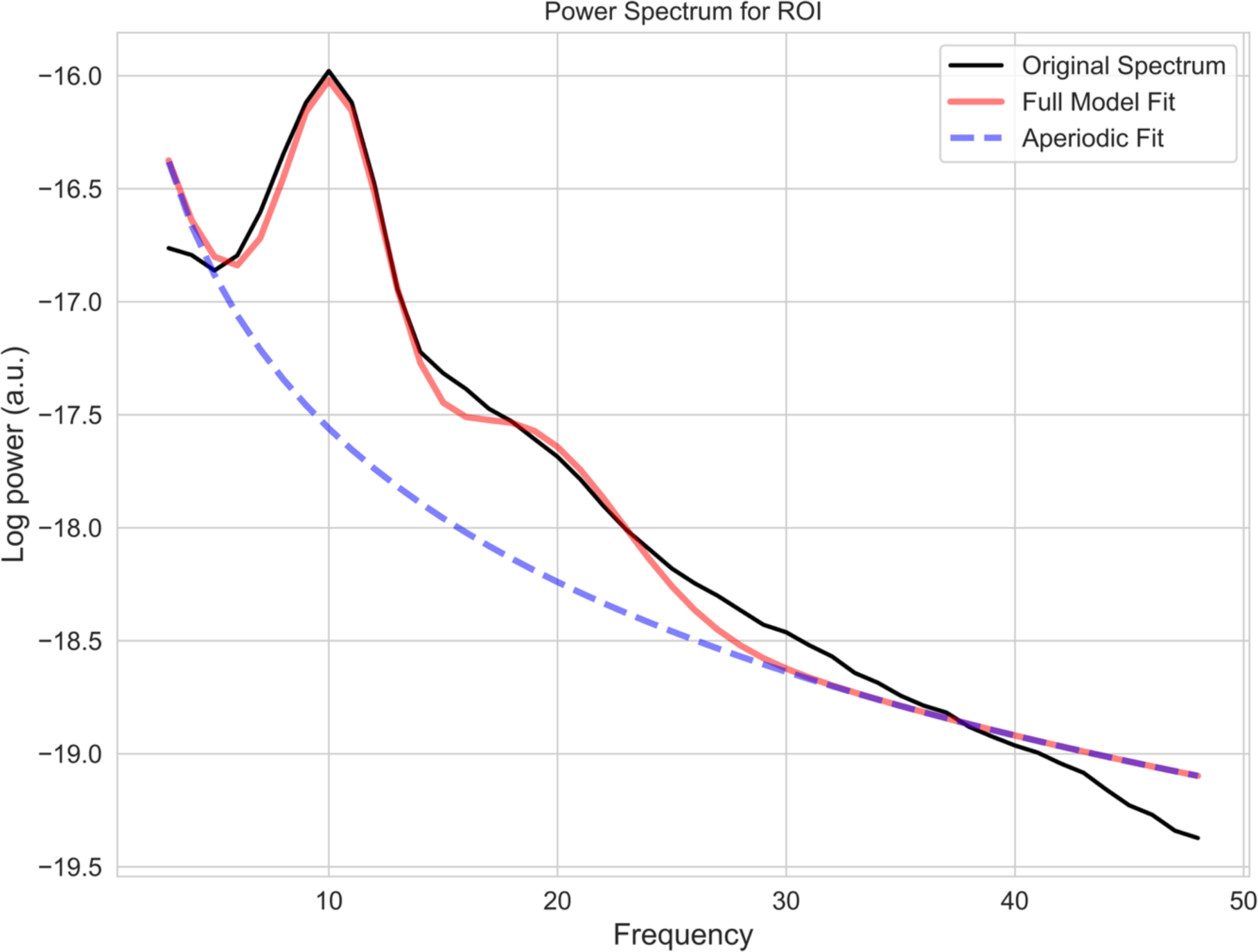
Example plot with fitted parameters (associated with participant ‘sub-032302’ right cuneus). The x-axis shows the frequency band in Hz units. The y-axis shows power in arbitrary units (a.u.) associated with resting-state activity in the eyes-closed condition at the right cuneus ROI (referred to the Desikan-Killiany atlas) for this participant.

**Figure 2.**
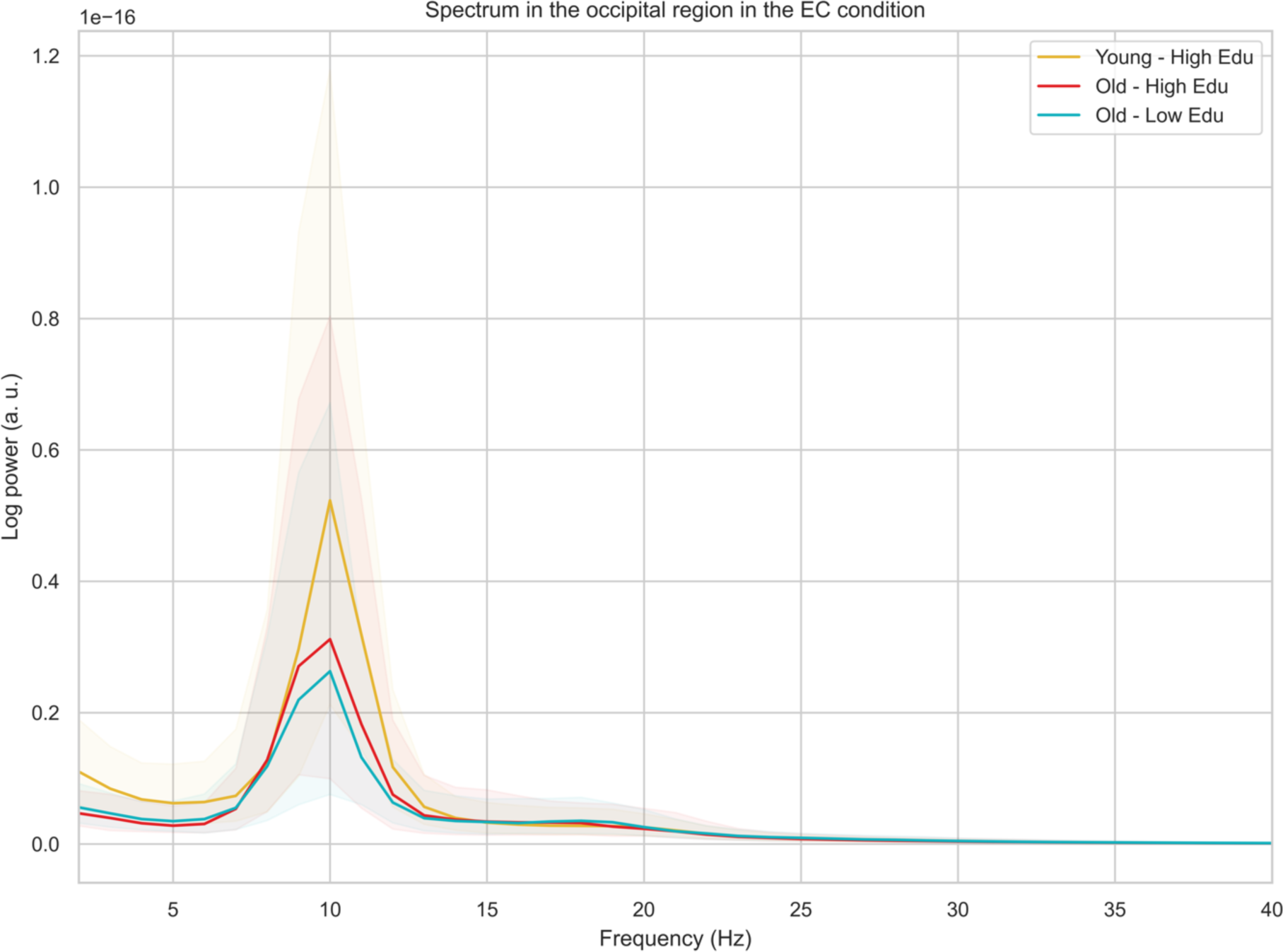
Age- and education-related differences in resting EEG power spectra in the occipital region, in the eyes closed condition. The plot shows a median for each group (Young – High Edu = younger adults with high education, Old – High Edu = older adults with high education; Old – Low Edu = older adults with low education) and a 50% percentile interval, ranging from the 25 to 75 percentiles.

**Figure 3.**
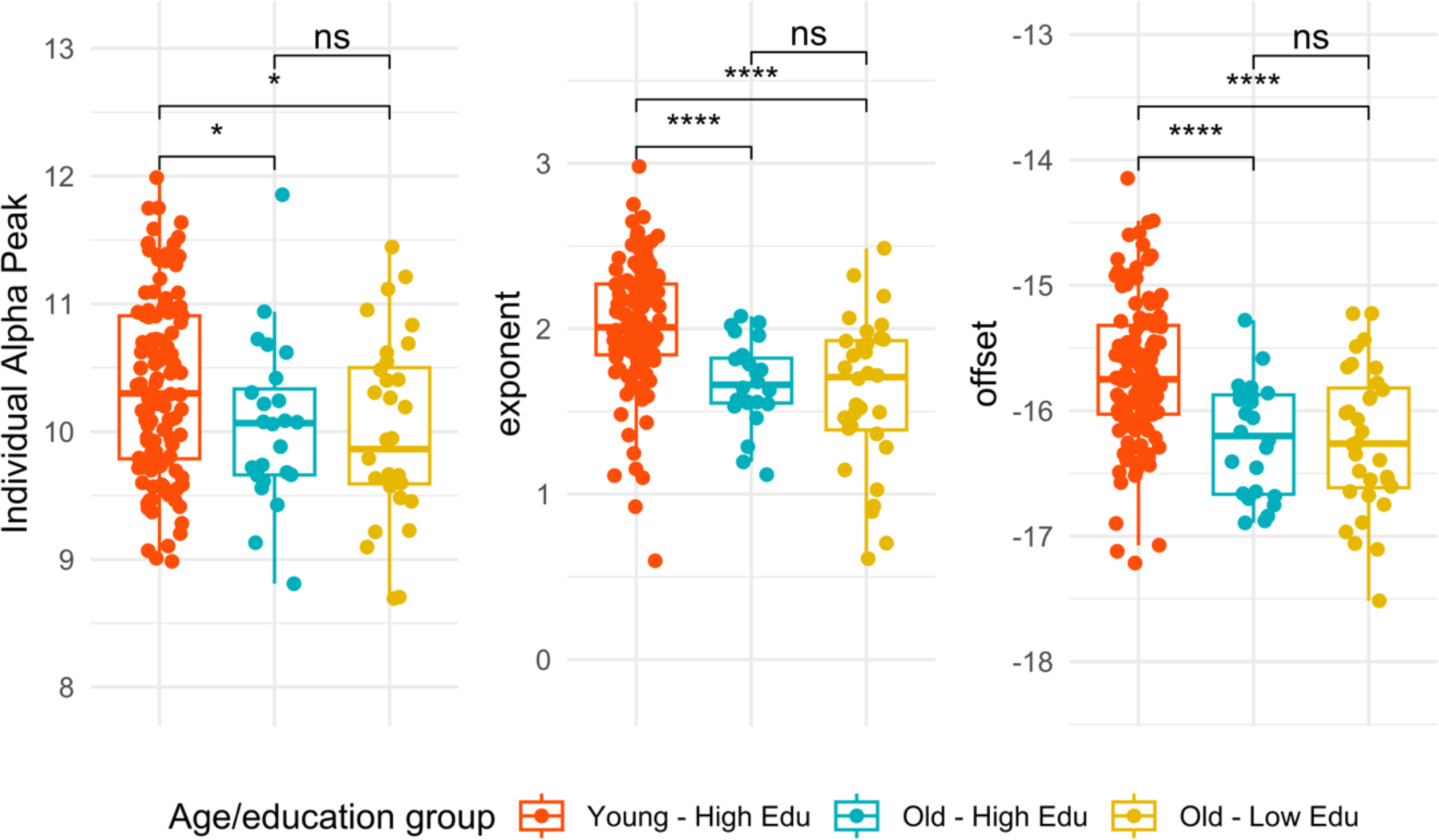
Age- and Education-related differences of the EEG components. The Individual Alpha Peak Frequency, the exponent, and the offset of each group (Young – High Edu = younger adults with high education, Old – High Edu = older adults with high education; Old – Low Edu = older adults with low education) are shown on the y-axis.

**Table 1.** Descriptive information about the sample and group comparisons based on participants’ age and education. Table depicts: Processing Speed alertness (PS_Allertness), Processing Speed Visual Attention (PS_Attention), Working Memory accuracy (MEM_WM) and Delayed Memory accuracy (MEM_Del); gray matter volume normalized (GM_Volume), Individual Alpha Peak Frequency (IAPF), Exponent, and Offset).

|  | Young Adults |  | Older Adults with high education |  | Older Adults with low education |  | Group comparison<br>Young – High Edu vs Old – Low Edu |  | Group comparison<br>Young – High Edu vs Old – Low Edu |  | Group comparison<br>Old – Low Edu vs Old – Low Edu |  |
| --- | --- | --- | --- | --- | --- | --- | --- | --- | --- | --- | --- | --- |
| | Range | M(SD) | Range | M (SD) | Range | M (SD) | $\chi^2$ | *p* | $\chi^2$ | *p* | $\chi^2$ | *p* |
| **PS_Allertness** | (-1.47 - 1.50) | -0.43 (0.56) | (-0.99 - 3.35) | 0.69 (1.07) | (-0.66 - 5.29) | 0.77 (1.07) | 23.15 | <0.001 | 30.44 | <0.001 | <0.01 | 0.92 |
| **PS_Visual Attention** | (-1.47 - 1.50) | -0.43 (0.56) | (-0.71 - 3.62) | 0.94 (1.07) | (-0.84 - 3.59) | 0.97 (1.07) | 42.84 | <0.001 | 42.11 | <0.001 | 0.08 | 0.77 |
| **MEM_WM** | (-2.83 - 0.80) | 0.24 (0.71) | (-3.27 - 0.80) | -0.04 (1.28) | (-3.72 - 0.80) | -0.92 (1.28) | 0.64 | 0.42 | 26.30 | <0.001 | 8.13 | <0.01 |
| **MEM_Del** | (-2.46 - 1.27) | 0.30 (0.84) | (-3.51 - 0.79) | -0.85 (0.97) | (-2.46 - 1.08) | -0.54 (0.97) | 24.16 | <0.001 | 23.20 | <0.001 | 0.98 | 0.32 |
| **GM_Volume** | (0.42 - 0.50) | 0.46 (0.01) | (0.36 - 0.44) | 0.41 (0.01) | (0.38 - 0.45) | 0.42 (0.01) | 56.63 | <0.001 | 65.30 | <0.001 | 2.07 | 0.14 |
| **IAPF** | (8.95 - 11.98) | 10.38 (0.68) | (8.81 - 11.85) | 10.04 (0.67) | (8.69 - 11.44) | 10.00 (0.67) | 3.96 | 0.04 | 5.46 | 0.01 | 0.17 | 0.67 |
| **Exponent** | (0.59 - 2.71) | 2.00 (0.36) | (1.11 - 2.07) | 1.67 (0.38) | (0.61 - 2.48) | 1.61 (0.38) | 22.19 | <0.001 | 20.77 | <0.001 | 0.21 | 0.64 |
| **Offset** | (-17.21 - 14.55) | -15.72 (0.54) | (-16.89 - 15.28) | -16.23 (0.51) | (-17.52 - 15.22) | -16.24 (0.51) | 18.33 | <0.001 | 18.61 | <0.001 | 0.02 | 0.88 |

### 3.2. EEG spectral parameters and cognitive performance

In the whole sample (179) a higher exponent and offset significantly predicted a better performance on the visual attention task (exp: *B* = -0.44, *p* < 0.01; offset: *B* = -0.32, *p* < 0.01). The exponent and the offset values predicted working memory capacity (exp: *B* = 0.37, *p* = 0.04; offset: *B* = 0.31, *p* = 0.01); see Section C of Supplementary Materials. Significant results emerged when considering the three groups for both the exponent and the offset, in the visual attention and working memory tasks. More specifically, compared to the group of young adults, where exponent and offset did not predict any variation in cognitive performance, older adults showed significant associations, different across educational levels. In the visual attention task, low educated older adults had a better (faster) performance at the higher aperiodic values (exp: *B* = -0.67, *p* = 0.04; offset: *B* = -0.56, *p*=0.03) while highly educated ones had a worse performance (slower) at the higher values of the exponent (exp: *B* = 1.41, *p*=0.02), a result confirmed post-hoc through slope comparisons, which confirmed a general age-related effect and also a significant difference between older adults with different education, at the third quartile of the exponent values (t=2.45; p = 0.03), see Figure 4.

**Figure 4.**
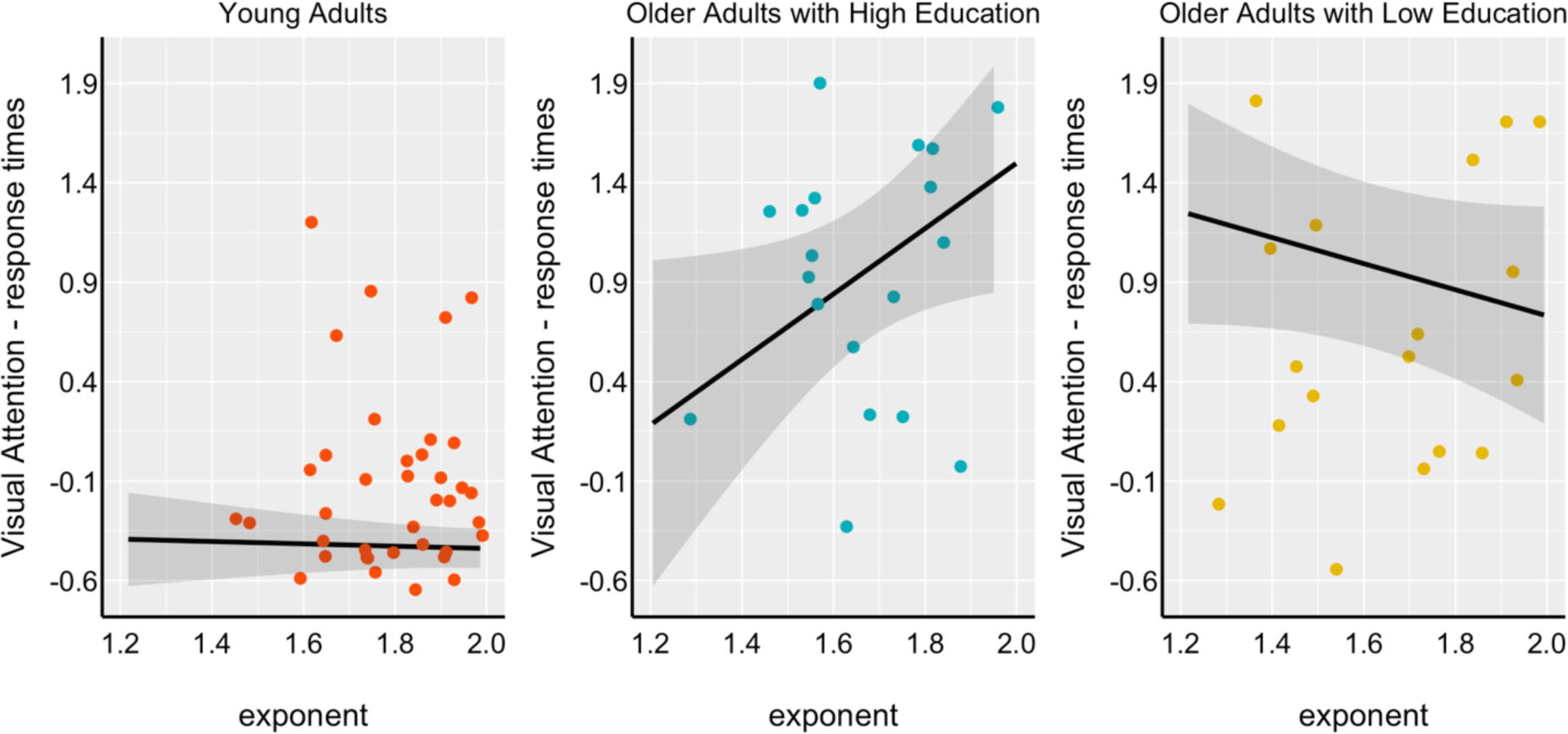
Relationship between exponent and response times in the visual attention task (processing speed). On the x-axis, the exponent value parameterized at the occipital level and in the Alpha band (8-12 Hz) is reported. On the y-axis, the z-scores associated with the Processing Speed response times: visual attention (i.e., “Trail Making Test-B”) is reported. The higher the exponent, the slower older adults with higher education, the faster the older adults with lower education. A non-significant relationship was shown between exponent and response times in the group of young adults.

Results showed that, also in the working memory task, older adults with high education had worse performance with increasing exponent values, as compared with those with low education (*B* = - 1.71, *p*=0.03). Post-hoc slope comparison showed no main education-related difference among older adults. Young adults and highly educated older adults did not differ between each other at different quartiles of exponent values (25%: t=1.16, p = 0.47; 50%: t=1.94, p = 0.12; 75%: t=2.12, p = 0.08); Figure S3 in Supplementary Materials and Tables S2-5 for more details.

## 4. DISCUSSION

The present study aimed to investigate the relationship between aperiodic activity and cognitive performance, by accounting for the level of education in older individuals, compared with a control group of highly educated younger adults (i.e., high neurocognitive efficiency).

The older adults showed less cognitive efficiency compared to young adults, across all tasks, which aligns with the well-established literature about cognitive decline in healthy aging ^41^. Upon stratifying older adults based on their education, the results indicated that those with higher education exhibited comparable performance to older adults with lower education, except for working memory performance. Older adults with higher education showed better working memory performance compared with older adults with lower education, which made the group of older adults with high education more similar to the young group, suggesting a potential role of education in mitigating age-related cognitive decline.

Consistent with previous evidence ^28,42,43^, the periodic and aperiodic components of EEG differentiated between young and older adults: older adults exhibited lower values across components, compared to younger adults. Concerning the periodic component of the EEG signal, IAPF showed the pattern of an age-related slowing, as confirmed in different studies ^44,45^. Several interpretations have been advocated to link periodic EEG components with aging. In addition to slowing with age, structural alterations in the brain have also been associated with the decline in power and peak frequency of alpha oscillations, particularly in older individuals ^10,46^. Additionally, the stability of IAPF over the life course reflects the preserved functionality of the central nervous system ^47^. Regarding cognition, IAPF has previously demonstrated a positive relationship with interference resolution in working memory performance, primarily observed in the temporal lobes^48^. This may indicate that IAPF plays a functional role in the ability to disregard and suppress interfering information. However, our study did not confirm the relationship between IAPF and working memory, suggesting that other factors, such as the specific nature of the task may mediate it.

In relation to the aperiodic EEG components, we found that both exponent and offset values significantly decreased with age. These results corroborate previous evidence suggesting that the aperiodic EEG component can serve as a neurophysiological marker of aging. Likewise, recent studies have revealed that aperiodic activity is influenced by various factors, including drugs ^49,50^ and level of arousal ^51^. However, the potential mediation of education, specifically its influence on the relationship between aperiodic components and cognitive performance, was unexplored in the literature.

In our study, education might help in interpreting the relationship between aperiodic component and performance on some tasks of visual attention and working memory, but not on a delayed memory task. The relation between aperiodic component and cognitive performance varied depending on education level, with a reversed pattern between exponent and cognitive performance in older adults across higher vs lower education. Older adults with lower education displayed a positive relationship between exponent and cognitive performance, while those with higher education exhibited the opposite trend. In this sense, the neurobiological bases of the aperiodic activity are still uncertain. However, evidence suggests that a flatter exponent (when the exponent approximates zero) may reflect an increase of asynchronous background neuronal firing, commonly called neural noise ^43^. Moreover, in non-linear systems like the brain, the notion of stochastic resonance proposes that information at the threshold level can be better processed within an optimal noise range than without noise ^52^. If different exponent values represent varying levels of neural noise, it is possible that noise also has different effects on performance according to the system. In older adults with lower education levels, higher exponents - corresponding to lower noise values - may contribute to better performance. On the other hand, older adults with higher education would exhibit the opposite pattern. In this latter group, higher exponents (lower noise values) would reduce performance efficiency. These two scenarios may depend on the fact that, according to the framework of stochastic resonance, there is no ideal level of noise and its effect on performance may not follow a linear pattern: it can vary based on the specific system and compensatory dynamics. Although such result may seem counterintuitive, a similar reversed pattern has been observed in a previous study that examined the relationship between mathematical achievement and glutamate concentrations. Glutamate has the effect of flattening the power spectrum, leading to exponent values closer to zero^49^. A study by ^53^ has demonstrated that the concentration of glutamate and the exponent levels could result in reversed cognitive performance outcomes depending on the participants’ age. Specifically, in their study, the concentration of glutamate (in the intraparietal sulcus) was negatively associated with mathematical achievement in younger participants, but it was positively associated with mathematical achievement in older participants. In our study, where the observed effects on cognitive performance cannot be influenced by age within the group of older adults, they could be associated with educational levels, which is a distinguishing factor between the two aging groups. Although tantalizing, this proposed relationship between exponent values, noise levels, and cognitive performance in older adults at different education level should be interpreted cautiously. Indeed, education may mediate the relationship between the aperiodic component and cognitive performance differently. For example, it may impact cognitive strategies, task engagement, and compensatory mechanisms, leading individuals with higher education to have a high cognitive performance independently of the specific neural mechanisms reflected by aperiodic EEG component. We did observe a relationship between the exponent and processing speed, in line with ^54^. Moreover, the study partially replicated what it was found in some previous studies where a relationship between exponent and working memory performance was identified ^16,28^.

Overall, our results cannot be interpreted as exhaustive; they should emphasize the importance of considering the aperiodic component of EEG signal as a marker of neurophysiological mechanisms that relates with performance, which can be mediated by different aspects. Our study, in particular, focused on education as one of these aspects. An important limitation is related to the fact that the LEMON database, despite having many advantages, did not have the optimal characteristics for the aims of this study. In particular, it included a cohort of participants with different age (whereas a longitudinal dataset would be more suited) and it included a different number of participants for each group. Future longitudinal studies with better stratification should explore the ontogenetic trajectory of the exponent, to further investigate its role in cognitive performance during aging. In the present study, the availability of age and education variables in a categorical form might have limited the assessment of neurobehavioral relationships and the potential use of finer analysis modeling.

Future studies may explore the intricate connection between EEG parameters and cognition, by encompassing a broader range of variables that could modulate such a relationship. Another relevant aspect is related to the nature of this study: although it could provide some insights into relationships among the variables, it cannot establish causation.

In summary, this study may open to future research on the modulatory role of education and other cognitive reserve proxies, in the complex relationship between aperiodic EEG component and cognitive efficiency in aging.

## 2. METHODS

### 2.1. Participants and Materials

All participants included in this study were taken from the “Leipzig Study for Mind-Body-Emotion Interactions’’ (LEMON; ^55^. The final sample consisted of N=179 individuals. Socio-demographic information like age and education was shared in bins ^55^, not continuous. Participants were grouped based on age (older vs. younger adults) and education (high vs. lower). Participants included in the young group (N=123) aged 20-35 years and had high education levels (12 years of *lyceum/gymnasium*), whereas the group of older adults (N=56), age range 60-77 years, was divided in two groups: one with high education (12 years of *lyceum/*gymnasium, N=24) and the other with low education (10 years of *technical high school/Realschule*, N=32). Participants with no availability of EEG data and those who were indicated as with alcohol abuse or dyslexia problems were not included in the final data sample. A small subgroup of young adults with low education was not included in the sample according to the study purpose (N=7); one participant resulted as outlier on both visual inspection of residuals and Cook’s threshold and it was removed. Data collection was performed upon the Declaration of Helsinki and approved by the local ethics committee ^55^; all participants provided written informed consent before data acquisition for the study.

### 2.2. Cognitive Assessment

Processing Speed and Memory capacity were investigated in relationship with periodic and aperiodic EEG components.

- Processing Speed included alertness and visual attention and it was assessed using two tasks: the Test of Attentional Performance ^56^ and the Trail Making Test ^57^. The former estimated alertness: i.e., participants were asked to respond, as fast as possible, to the appearance of a visual stimulus on the screen. The TMT estimated visual attention w, i.e., participants were asked to connect as fast as possible a series of visual stimuli, alternatively with a definition order: numerical and alphabetical orders.

- Memory included working and delayed memory tasks; it was assessed with a working memory task (WM_TAP,^56^ and the California Verbal Learning Task (CVLT, ^58^. For the working memory task, participants had to simultaneously provide a response only when a given stimulus was equal to the second last one perceived in the series, while keeping track of a series of different stimuli. In the delayed memory task, participants were asked to retain and correctly recall a series of 16 words belonging to their vocabulary.

### 2.3 Neural variables

#### 2.3.1. Gray Matter Volume

A 3 Tesla scanner (MAGNETOM Verio, Siemens Healthcare GmbH, Erlangen, Germany) with a 32-channel head coil was used to conduct Magnetic Resonance Imaging (MRI); ^55^. The pre-processing pipeline included a series of steps: a) re-orientating images to the standard (MNI) template, b) bias field correction, c) registration to the MNI template using both linear (FLIRT) and nonlinear (FNIRT) registration tools, and d) brain extraction. Brain tissues were segmented using FMRIB’s Automated Segmentation Tool (FAST) that allowed extracting measures of total Gray Matter, White Matter and Cerebrospinal Fluid. Brain tissues were visually inspected by a trained neuroscientist (NF) to ensure an accurate segmentation.

#### 2.3.2. EEG preprocessing and source reconstruction

A 16-min rs-EEG was recorded with a BrainAmp MR plus amplifier in an electrically shielded and sound-attenuated EEG booth using 62-channel (61 scalp electrodes plus 1 electrode recording the VEOG below the right eye) active ActiCAP electrodes (Brain Products GmbH, Gilching, Germany; international standard 10–20 localization system, and referenced to FCz). EEG was recorded with a band-pass filter between 0.015 Hz and 1 kHz and digitized with a sampling rate of 2500 Hz. Raw EEG data were down-sampled from 2500 Hz to 250 Hz and band-pass filtered within 1-45 Hz. Outlier channels were rejected after visual inspection for frequent jumps/shifts in voltage and poor signal quality. Data intervals containing extreme peak-to-peak deflections or large bursts of high-frequency activity were identified by visual inspection and removed. Independent component analysis (ICA) was performed using the Infomax algorithm (runica function from MATLAB). On pre-processed files, source reconstruction was run by using a standard head model. A 3-shell boundary element model was constructed via Brainstorm ^59^. The default current density maps were normalized through Standardized LOw Resolution brain Electromagnetic TomogrAphy approach (sLORETA, ^60^. Welch’s method was used to calculate the power spectrum at the level of the reconstructed sources. Window-overlap was 50%. Due to the small number of EEG channels, we grouped cortical vertices into major regions (ROIs), aggregated accordingly to Desikan-Killiany atlas by following a similar approach ^48^.

#### 2.3.3. Periodic and aperiodic components of the power spectral density

The specparam algorithm (version 1.0.0 ^16^ was used to parametrise power spectra of ROIs. In specparam algorithm, the power spectrum *PSD* is modeled as a combination of aperiodic component *AP* and a sum of N oscillatory peak modeled with a Gaussian:

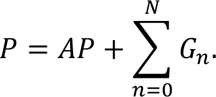

The component *AP*(*f*) for frequency *f* is expressed by the formula:

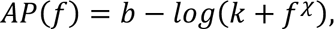

where *b* is the broadband offset, *χ* is the exponent and *k* is the knee parameter, controlling the “bend”, When *k* = 0 this the component *AP* will be a line fitted in the log-log space (this is later referred as a fixed mode). in this case, the slope of the line *a* in log-log space is directly related to exponent *χ*, *χ* = −*a* ^16^. The output of algorithm for estimated peaks are the mean of the Gaussian *G_n_* for the center frequency of the peak, aperiodic-adjusted power (the distance between the peak of the Gaussian and the aperiodic fit at this frequency) and bandwidth as 2 *SD* of the fitted Gaussian. In the current analysis, power spectra were parameterized across the frequency range 3 to 48 Hz (the maximal frequency range to avoid the line noise frequency) using the “fixed” mode (Figure 1). Additional algorithm settings were set as: peak width limits: [2.5 8]; max number of peaks: 6; minimum peak height: 0.5; peak threshold: 2. All the parameters describing identified peaks, offset, exponent and the parameters describing how well the model was fit were extracted.

The parameters were extracted for every PSDs in the Eyes Closed condition, then ROIs that belong to the occipital regions were selected (that is where dominant activity in alpha is expected to exhibit age-related patterns; ^44,45,47^; thus the parameters from all ROIs belonging to this region were averaged. The choice of parameters gave a median goodness-of-fit measure of *r*^&^ = 0.981, IQR= [0.971,0.978] across all regions within the occipital lobe. Model fits were not statistically different between the two groups: YA median r2 = 0.983, IQR= [0.974,0.988]; OA median r2 = 0.976, IQR= [0.962, 0.984]. Thus, although other processing parameters could have been chosen, we achieved suitable spectral parameterization across participants and regions. Compared to previous studies using the specparam algorithm, where frequency range varies, many used 40 Hz as the upper frequency range ^23,48,61^. For sake of clarity, using 48 Hz as upper frequency is also a widely adopted option ^62,63^, and stays within the recommendations of the algorithm’s authors. All participants showed a discernible alpha peak in the PSD (see an example in Figure 1). Individual alpha peak frequency values per subject were created from periodic components fitted by algorithm in the alpha range (8-12 Hz), but instead of averaging several of them per subject per ROI only the one that had the highest value of PW was taken per ROI and then averaged across ROIs within the occipital region. Figure 2 offers a qualitative overview of the differences in EEG power spectra among our groups. The figure visually confirms differences in IAPF (Individual Alpha Peak Frequency) values between the young and old populations.

### 2.4. Statistical Analyses

Analyses were performed with the R software ^64^. Shapiro-Wilk tests on each dependent variable, inspection of residuals, and Kolmogorov-Smirnov analysis were performed to build up the regression models. Results indicated General Linear regression Models as suitable; they included processing speed and accuracy scores on cognitive tests as dependent variables, the variable group as a factor: older adults with high education vs. older adults with lower education vs. young adults (all high education). Other continuous predictors of interest were the periodic and the aperiodic EEG components: IAPF, exponent, and offset values. Sex and normalized gray matter volume were accounted for in all regression models. A simplified syntax of the R linear models is reported below:

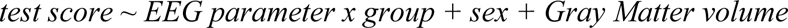

Sex and Gray matter volume were added as covariates as they were two relevant variables that could also be associated with cognitive performance. Power analysis revealed a statistical power > 0.95 indicating the ability of the model to detect a significant effects, based on a significance level (α) of 0.05 and an estimated effect size f² of 0.35.

