## Supplementary File for "Aperiodic component of EEG power spectrum and cognitive performance in aging: the role of education"

### TABLE OF CONTENTS

***SECTION A: Group-differences*3**

***SECTION B: Regression Analysis*5**

### SECTION A: GROUP-DIFFERENCES

Younger adults were overall more efficient in performing cognitive tasks than older adults with both high and lower education, except in the case of working memory tasks, where older adults with higher education did not differ from younger adults.

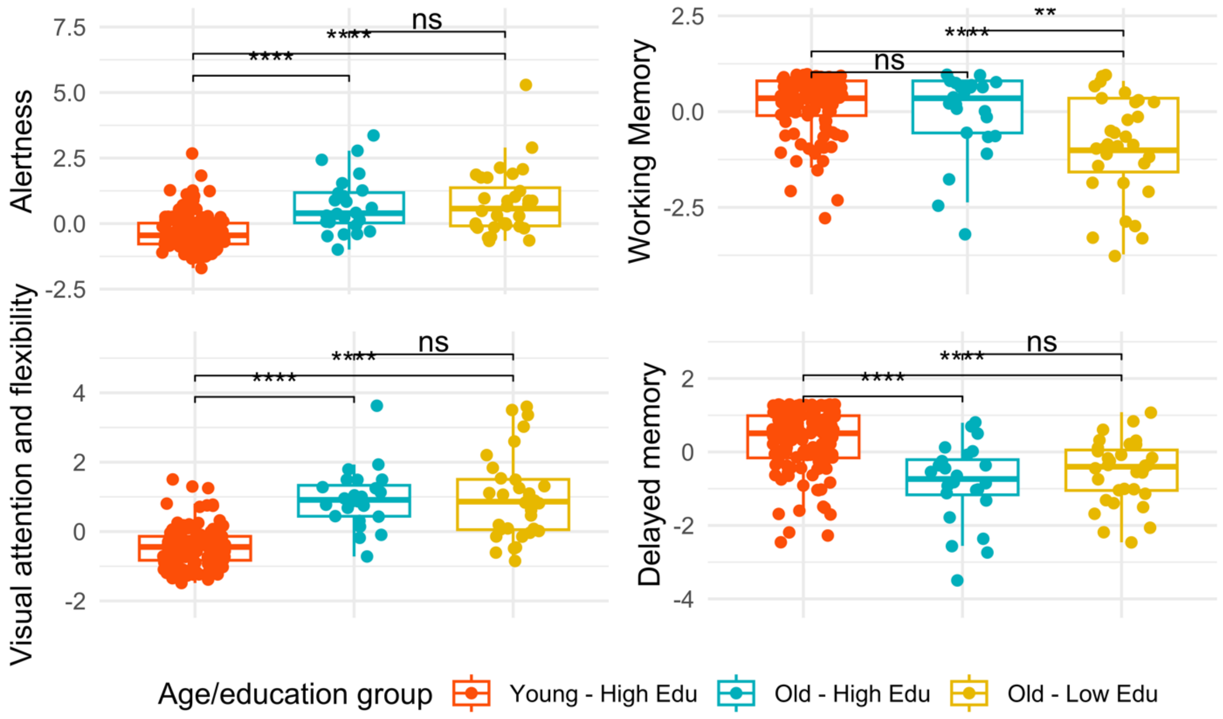


**Figure S1. Group Comparisons of cognitive variables.** The panel shows, on the upper left-hand side the group-differences associated with the Alertness processing speed, on the upper right-hand side the group differences associated with the Working Memory accuracy; on the lower left-hand side, the differences associated with Visual Attention processing speed; on the lower right-hand side with Delayed Memory accuracy. Significance is set with $p<0.05$.

| | Young Adults (38 F) | | Older Adults (26 F) | | Young - Old |
|---|---|---|---|---|---|
| | Range | M(SD) | Range | M (SD) | *p* |
| **PS_Attention** | (-1.47 - 1.50) | -0.43 (0.56) | (-0.84 - 3.63) | 0.97 (1.07) | <0.001 |
| **MEM_WM** | (-2.83 - 0.80) | 0.24 (0.71) | (-3.73 - 0.80) | -0.55 (1.28) | <0.001 |
| **MEM_Del** | (-2.46 - 1.27) | 0.30 (0.84) | (-3.52 - 1.08) | -0.68 (0.97) | <0.001 |
| **GM_Volume** | (0.42 - 0.50) | 0.46 (0.01) | (0.36 - 0.45) | 0.41 (0.01) | <0.001 |
| **IAPF** | (8.95 - 11.98) | 10.38 (0.68) | (8.69 - 11.85) | 10.03 (0.67) | <0.01 |
| **Exponent** | (0.59 - 2.71) | 2.00 (0.36) | (0.61 - 2.48) | 1.63 (0.38) | <0.001 |
| **Offset** | (-17.21 - 14.55) | -15.72 (0.54) | (-17.52 - 15.22) | -16.24 (0.51) | <0.001 |

**Table S1. Descriptive information about the sample, based on participants' age.** Age-related group comparisons on Processing Speed alertness (PS_Allertness), Processing Speed Visual Attention (PS_Attention), Working Memory accuracy (MEM_WM) and Delayed Memory accuracy (MEM_Del). Brain measures reported are gray matter volume normalized (GM_Volume), Individual Alpha Peak Frequency (IAPF), Exponent, and Offset.

### SECTION B: Regression analyses

| Processing Speed: Alertness | | Estimate | Std. Error | t-value | Pr(>\|t\|) |
|---|---|---|---|---|---|
| IAPF | (Intercept) | -0,27 | 2,82 | -0,10 | 0,92 |
| | GM_Normalised | -3,23 | 3,95 | -0,82 | 0,42 |
| | sexM | -0,11 | 0,15 | -0,77 | 0,44 |
| | Young – High Edu | 1,20 | 2,53 | 0,47 | 0,64 |
| | Old – High Edu | 3,17 | 3,66 | 0,87 | 0,39 |
| | mean_peak_alpha | 0,25 | 0,22 | 1,11 | 0,27 |
| | Young – High Edu: IAPF | -0,22 | 0,25 | -0,86 | 0,39 |
| | Old – High Edu: IAPF | -0,32 | 0,36 | -0,89 | 0,38 |
| exponent | (Intercept) | 3,23 | 1,82 | 1,77 | 0,08 |
| | GM_Normalised | -4,55 | 3,97 | -1,15 | 0,25 |
| | sexM | -0,10 | 0,14 | -0,70 | 0,48 |
| | Young – High Edu | -1,20 | 0,73 | -1,64 | 0,10 |
| | Old – High Edu | -2,56 | 1,36 | -1,89 | 0,06 |
| | mean_exponent | -0,30 | 0,34 | -0,89 | 0,38 |
| | Young – High Edu: exponent | 0,21 | 0,40 | 0,53 | 0,59 |
| | Old – High Edu: exponent | 1,49 | 0,80 | 1,86 | 0,06 |
| offset | (Intercept) | -1,32 | 4,66 | -0,28 | 0,78 |
| | GM_Normalised | -3,62 | 3,99 | -0,91 | 0,37 |
| | sexM | -0,14 | 0,15 | -0,93 | 0,36 |
| | Young – High Edu | 0,89 | 5,00 | 0,18 | 0,86 |
| | Old – High Edu | 4,67 | 8,02 | 0,58 | 0,56 |
| | mean_offset | -0,23 | 0,28 | -0,82 | 0,41 |

| **Processing Speed: Alertness** | | Estimate | Std. Error | t-value | Pr(>\|t\|) |
|---|---|---|---|---|---|
| | Young – High Edu: offset | 0,11 | 0,31 | 0,35 | 0,73 |
| | Old – High Edu: offset | 0,29 | 0,49 | 0,59 | 0,56 |

**Table S2. Summary output of the Regression model with alertness response times as dependent variable.** The table displays, in the first column, Individual Alpha Peak Frequency (IAPF), exponent, and offset; in the second column, the terms of the mode. The third, fourth, and fifth columns refer to the B (estimate) values, the relative standard error, the t-value associated with the fitted term, and the p-value.

| **Processing Speed: Visual Attention** | | Estimate | Std. Error | t-value | Pr(>\|t\|) |
|---|---|---|---|---|---|
| IAPF | (Intercept) | 2,16 | 2,46 | 0,88 | 0,38 |
| | GM_Normalised | 0,52 | 3,44 | 0,15 | 0,88 |
| | sexM | 0,21 | 0,13 | 1,61 | 0,11 |
| | Young – High Edu H | -2,69 | 2,21 | -1,22 | 0,23 |
| | Old – High Edu | -0,53 | 3,20 | -0,17 | 0,87 |
| | mean_peak_alpha | -0,15 | 0,19 | -0,76 | 0,45 |
| | Young – High Edu: IAPF | 0,12 | 0,22 | 0,55 | 0,58 |
| | Old – High Edu: IAPF | 0,04 | 0,32 | 0,14 | 0,89 |
| exponent | (Intercept) | 0,07 | 1,64 | 0,04 | 0,97 |
| | GM_Normalised | -1,06 | 3,38 | -0,31 | 0,75 |
| | sexM | 0,24 | 0,12 | 1,95 | 0,05 |
| | Young – High Edu | 2,50 | 0,62 | 4,01 | <0,001 |
| | Old – High Edu | -1,06 | 1,14 | -0,93 | 0,35 |
| | mean_exponent | -0,09 | 0,18 | -0,50 | 0,62 |
| | Young – High Edu: exponent | -0,68 | 0,34 | -1,98 | 0,05 |
| | Old – High Edu: exponent | 1,41 | 0,64 | 2,21 | 0,03 |
| offset | (Intercept) | -1,42 | 2,50 | -0,57 | 0,57 |
| | GM_Normalised | -0,33 | 3,41 | -0,10 | 0,92 |
| | sexM | 0,20 | 0,12 | 1,57 | 0,12 |
| | Young – High Edu | -7,72 | 4,28 | -1,81 | 0,07 |
| | Old – High Edu | 6,09 | 5,95 | 1,03 | 0,31 |
| | mean_offset | -0,06 | 0,12 | -0,53 | 0,60 |
| | Young – High Edu: offset | -0,56 | 0,26 | -2,13 | 0,03 |

| **Processing Speed: Visual Attention** | | Estimate | Std. Error | t-value | Pr(>\|t\|) |
|---|---|---|---|---|---|
| | Old – High Edu: offset | 0,29 | 0,37 | 0,80 | 0,43 |

**Table S3. Summary output of the Regression model with visual attention response times as dependent variable.** The table displays, in the first column, Individual Alpha Peak Frequency (IAPF), exponent, and offset; in the second column, the terms of the mode. The third, fourth, and fifth columns refer to the B (estimate) values, the relative standard error, the t-value associated with the fitted term, and the p-value.

| Working Memory | | Estimate | Std. Error | t-value | Pr(>\|t\|) |
|---|---|---|---|---|---|
| IAPF | (Intercept) | 2,44 | 2,34 | 1,04 | 0,30 |
| | GM_Normalised | -2,90 | 4,09 | -0,71 | 0,48 |
| | sexM | 0,12 | 0,15 | 0,79 | 0,43 |
| | Young – High Edu | -1,34 | 2,63 | -0,51 | 0,61 |
| | Old – High Edu | -3,35 | 3,26 | -1,03 | 0,31 |
| | mean_peak_alpha | -0,09 | 0,12 | -0,75 | 0,46 |
| | Young – High Edu: IAPF | 0,00 | 0,26 | 0,02 | 0,99 |
| | Old – High Edu: IAPF | 0,29 | 0,32 | 0,89 | 0,38 |
| exponent | (Intercept) | -1,36 | 1,88 | -0,72 | 0,47 |
| | GM_Normalised | -1,41 | 4,09 | -0,35 | 0,73 |
| | sexM | 0,10 | 0,15 | 0,66 | 0,51 |
| | Young – High Edu | 1,90 | 0,76 | 2,52 | 0,01 |
| | Old – High Edu | 3,68 | 1,40 | 2,63 | 0,01 |
| | mean_exponent | 0,61 | 0,35 | 1,74 | 0,08 |
| | Young – High Edu: exponent | -0,47 | 0,42 | -1,13 | 0,26 |
| | Old – High Edu: exponent | -1,72 | 0,82 | -2,09 | 0,04 |
| offset | (Intercept) | 8,40 | 4,77 | 1,76 | 0,08 |
| | GM_Normalised | -1,97 | 4,09 | -0,48 | 0,63 |
| | sexM | 0,15 | 0,15 | 0,99 | 0,32 |
| | Young – High Edu | -4,40 | 5,12 | -0,86 | 0,39 |
| | Old – High Edu | -11,79 | 8,21 | -1,44 | 0,15 |
| | mean_offset | 0,53 | 0,28 | 1,87 | 0,06 |
| | Young – High Edu: offset | -0,34 | 0,32 | -1,08 | 0,28 |

| **Working Memory** | | Estimate | Std. Error | t-value | Pr(>\|t\|) |
|---|---|---|---|---|---|
| | Old – High Edu: offset | -0,78 | 0,51 | -1,53 | 0,13 |

**Table S4. The summary output of the Regression model with working memory accuracy as a dependent variable.** The table displays, in the first column, Individual Alpha Peak Frequency (IAPF), exponent, and offset; in the second column, the terms of the mode. The third, fourth, and fifth columns refer to the B (estimate) values, the relative standard error, the t-value associated with the fitted term, and the p-value.

| Delayed Memory recall | | Estimate | Std. Error | t-value | Pr(>\|t\|) |
|---|---|---|---|---|---|
| IAPF | (Intercept) | -0,55 | 3,92 | -0,14 | 0,89 |
| | GM_Normalised | -0,52 | 0,15 | -3,53 | < 0.001 |
| | sexM | 0,84 | 2,52 | 0,33 | 0,74 |
| | Young – High Edu | 0,79 | 3,64 | 0,22 | 0,83 |
| | Old – High Edu | -0,05 | 0,22 | -0,22 | 0,83 |
| | mean_peak_alpha | 0,02 | 0,25 | 0,08 | 0,94 |
| | Young – High Edu: IAPF | -0,09 | 0,36 | -0,26 | 0,79 |
| | Old – High Edu: IAPF | | | | |
| exponent | (Intercept) | -0,23 | 1,82 | -0,13 | 0,90 |
| | GM_Normalised | -0,77 | 3,97 | -0,19 | 0,85 |
| | sexM | -0,50 | 0,14 | -3,45 | < 0.001 |
| | Young – High Edu | 1,25 | 0,73 | 1,71 | 0,09 |
| | Old – High Edu | -0,81 | 1,36 | -0,60 | 0,55 |
| | mean_exponent | 0,13 | 0,34 | 0,37 | 0,71 |
| | Young – High Edu: exponent | -0,14 | 0,40 | -0,34 | 0,73 |
| | Old – High Edu: exponent | 0,38 | 0,80 | 0,48 | 0,63 |
| | (Intercept) | -0,01 | 3,96 | 0,00 | 1,00 |
| | GM_Normalised | -0,49 | 0,14 | -3,39 | < 0.001 |
| | sexM | -2,06 | 4,96 | -0,42 | 0,68 |
| | Young – High Edu | -6,43 | 7,95 | -0,81 | 0,42 |
| | Old – High Edu | 0,23 | 0,27 | 0,83 | 0,41 |
| | mean_offset | -0,19 | 0,31 | -0,61 | 0,54 |
| | Young – High Edu: offset | -0,39 | 0,49 | -0,79 | 0,43 |

| **Delayed Memory recall** | | Estimate | Std. Error | t-value | Pr(>\|t\|) |
|---|---|---|---|---|---|
| | Old – High Edu: offset | -0,55 | 3,92 | -0,14 | 0,89 |

**Table S5. Summary output of the Regression model with delayed memory accuracy times as dependent variable.** The table displays, in the first column, Individual Alpha Peak Frequency (IAPF), exponent, and offset; in the second column, the terms of the mode. The third, fourth, and fifth columns refer to the B (estimate) values, the relative standard error, the t-value associated with the fitted term, and the p-value.

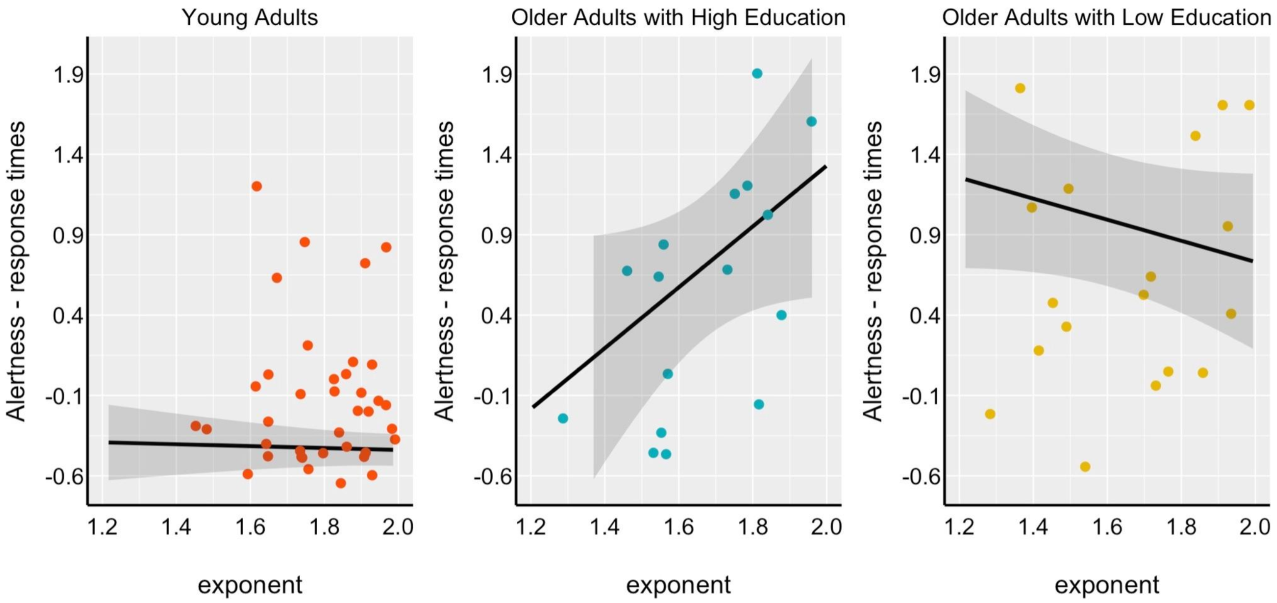


**Figure S2. Relationship between exponent and response times in the alertness task.** On the x-axis, the exponent value is parameterized at the occipital level in the Alpha band (8-12 Hz). On the y-axis, the z-scores associated with the Processing Speed response times: alertness (i.e., "TAP", Test of Attention Performance). The higher the exponent, the slower older adults with higher education, the faster the older adults with lower education. A non-significant relationship was shown between exponent and response times in the group of young adults.

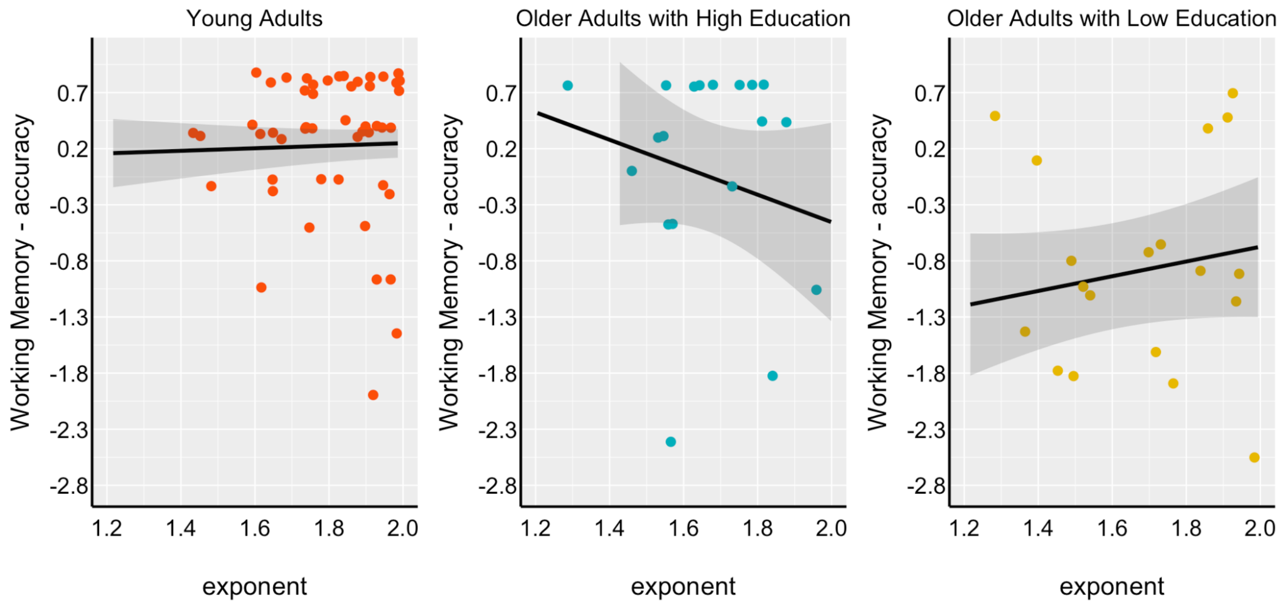


**Figure S3. Relationship between exponent and accuracy in the working memory task.** On the x-axis, the exponent value is parameterized at the occipital level in the Alpha band (8-12 Hz). On the y-axis, the z-scores are associated with Working Memory. No significant difference was shown across groups in the relationship between exponent and working memory task; however, a slightly stronger relationship can be found between these two variables in older adults with higher education.

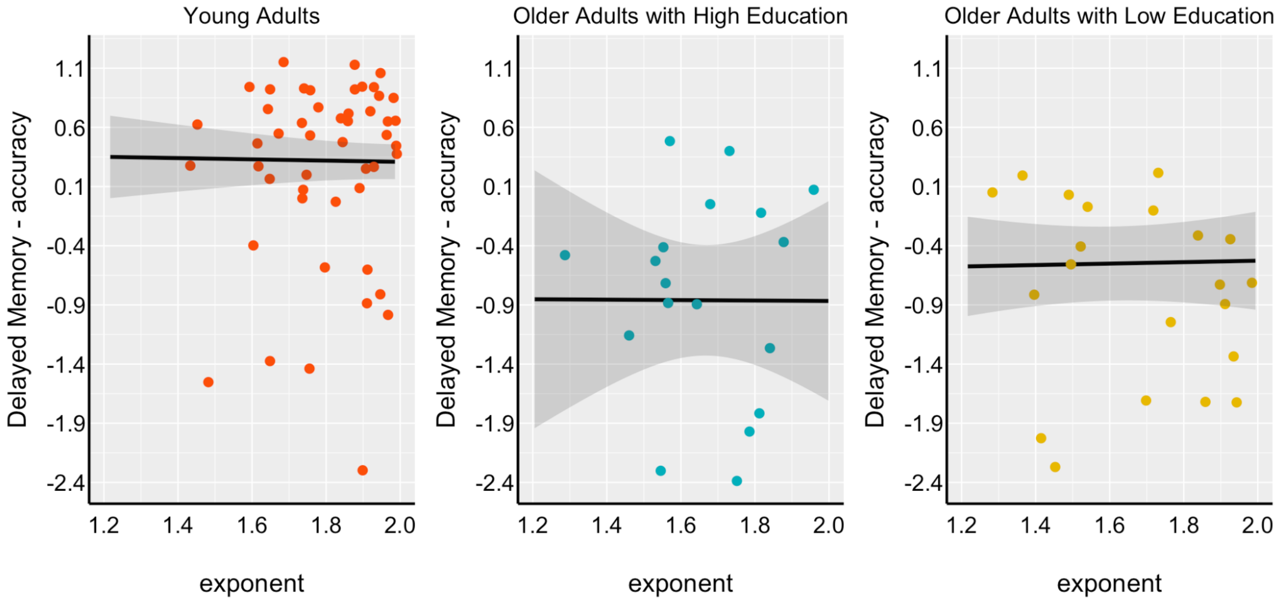


**Figure S4.** Relationship between exponent and accuracy in the delayed memory task. On the x-axis, the exponent value is parameterized at the occipital level in the Alpha band (8-12 Hz). On the y-axis, the z-scores are associated with Delayed Memory. No significant difference was shown across groups in the relationship between the exponent and the working memory task.
